# Evolution of fungal community associated with ready-to-eat pineapple during storage under different temperature conditions

**DOI:** 10.1101/2020.09.24.310573

**Authors:** Evanthia Manthou, Gwendoline Coeuret, Stephane Chaillou, George-John E. Nychas

## Abstract

The international market of fresh-cut products has witnessed dramatic growth in recent years, stimulated by consumer’s demand for healthy, nutritious and convenient foods. One of the main challenging issues for the quality and safety of these products is the potential microbial spoilage that can significantly reduce their shelf-life. The complete identification of fresh-cut product microbiota together with the evaluation of environmental factors impact on microbial composition is of primary importance. We therefore assessed the fungal communities associated with the spoilage of ready-to-eat (RTE) pineapple using a metagenetic amplicon sequencing approach, based on the ITS2 region. Our results revealed a significant variability on fungal species composition between the different batches of RTE pineapple. The initial microbiota composition was the main influencing factor and determined the progress of spoilage. Temperature and storage time were the secondary factors influencing spoilage and their impact was depending on the initial prevalent fungal species, which showed different responses to the various modifications. Our results strongly suggest that further large-scale sampling of RTE pineapple production should be conducted in order to assess the full biodiversity range of fungal community involved in the spoilage process and for unravelling the impact of important environmental factors shaping the initial microbiota.

## 1. Introduction

Fresh-cut market has grown dramatically in recent years, as a result of changes on consumers’ attitude. Ready-to-eat (RTE) fruits and vegetables fulfil the growing demand for healthy, convenient and minimally processed food products (Gorni et al., 2015; Qadri et al., 2015). However, the quality and safety assurance of these new types of fresh products is a major challenge for the fresh-cut industry and requires full involvement and increasing investigations of food scientists (Mederos et al., 2020).

Fresh-cut fruits and vegetables products have a limited shelf life due to accelerated physiological and biochemical changes occurring during their processing and storage (Zhang et al., 2014; Torri et al., 2010; Di Egidio et al., 2009). Indeed, the processing treatments render the products more prone to spoilage microorganisms, as well as micro-organisms of public health significance (Qadri et al., 2015; Leff and Fierer, 2013). Various studies underline the presence of phytopathogens and human pathogens, but also microorganisms with antagonistic properties against these pathogens, which have a significant influence on human health and products’ quality (Gorni et al., 2015). Therefore, a better insight into the microbial community and its potential interactions in food associated matrices is required to provide safe and high-quality food (Cao et al., 2017; Juste et al., 2008).

So far culture-dependent methods have been the gold standards in food microbiology, since they have led to the description of a number of habitats. However, they are extremely biased in their ability to unravel the microbial communities of complex matrices associated with food or environmental samples (Edet et al., 2017; Zhou et al., 2015; Ercolini et al., 2013; Juste et al., 2008). On the other hand, the development of next generation sequencing (NGS) techniques has enabled researchers to study food microbial ecology from broader and deeper perspectives. Recently, metagenetic and metagenomic approaches have resulted in improved understanding of a microbiota by providing a species- and strain-level characterization (Cauchie et al., 2020; Poirier et al., 2018; Abdelfattah et al., 2018; Abdelfattah et al., 2016). Most NGS related food microbiota studies have focused on fermented foods of different origins and to a lower extend on fresh meat and seafood products. NGS microbiota studies concerning fruits and vegetables are more limited and mainly focused on epiphytic microbial community (EFSA Panel on Biological Hazards, 2020; Angeli et al., 2019; Tatsika et al., 2019; Saminathan et al., 2018; Söderqvist et al., 2017; Abdelfattah et al., 2016; Dees et al., 2014; Jackson et al., 2013; Rastogi et al., 2012; Lopez-Velasco et al., 2010). Pineapple (*Ananas comosus*) is one of the most popular tropical fruit worldwide and it is commonly found in fresh-cut market. However, little have been reported about the associated microbial community and its response in various environmental factors (Dos Santos Souza et al., 2019; Di Cagno et al., 2010; Montero-Calderon et al., 2008). According to the limited studies based on the pineapple diversity, fungi have the leading role in fresh-cut pineapple’s spoilage. The fungal species, even the prevalent ones, reported in pineapple differ between the studies (Leneveu-Jevrin et al., 2020; Chanprasartsuk et al., 2010; Di Cagno et al., 2010; Tournas et al., 2006). Interestingly, the present literature is based on earlier generation molecular methods combined largely with culture-dependent and recently with culture-independent techniques.

In the present study, we used a metagenetic amplicon sequencing approach, based on the ITS2 region, with the objective to assess the fungal communities associated with the spoilage of RTE pineapple. To our knowledge, metagenetic analysis has never been applied to the pineapple microbiota. Therefore, this work sheds light on the variability of RTE pineapple’s spoilage microbiota and on how it is changing during shelf life with the influence of the temperature used during storage.

## 2. Materials and Methods

### 2.1. Sample preparation and storage conditions

Four batches of fresh-cut pineapple were supplied by a local manufacturer in Athens. The pineapple was packed in PVC trays each of one containing 220 g of fruit. The trays were transported to the laboratory within 24 hours from their production and stored at three different isothermal temperatures, 4, 8, 12°C and under dynamic temperature conditions (8 h at 4°C, 8 h at 8°C and 8 h at 12°C) in high precision (±0.5) incubators (MIR-153, Sanyo Electric Co., Osaka, Japan). The incubation temperature was recorded every 15 minutes using electronic temperature devices (COX TRACER®, Cox Technologies Inc., Belmont, NC, USA). The first sampling was conducted at the time of pineapple arrival to the laboratory and also at 38 h, 72 h, 134 h and 230 h of storage for 4, 8°C and the dynamic conditions. The final sampling point for 12°C was 134 h.

### 2.2. Microbial analysis and pH measurements

The samples (25 g of pineapple) were aseptically transferred into a sterile Stomacher bag, (Seward Medical, London, UK), diluted with 225 ml of Ringer buffer solution (Lab M Limited, Lanchashire, UK) and homogenized for 60 s at 230 rpm in a stomacher device (Lab Blender 400, Seward Medical, London, UK). The appropriate decimal progressive dilutions were prepared and the following microbial determinations were performed: total mesophilic microbial populations (total viable counts, TVC) by the spread method on tryptic glycose yeast agar (Plate Count Agar, Biolife, Milan, Italy), after incubation of plates at 25°C for 72 h; yeast and moulds by the spread method on rose bengal chloramphenicol agar (RBC, Lab M Limited) and incubation at 25°C for 3-5 days; *Pseudomonas* spp. by spread method on pseudomonas agar base with selective supplement cephalothin-fucidin-cetrimide (CFC, Lab M Limited) at 25°C for 48 h; lactic acid bacteria by pour method on de Man, Rogosa and Sharpe agar (MRS, Biolife) at 30°C for 72 h; and bacteria of the Enterobacteriaceae family by pour method on violet red bile glucose agar (VRBG, Biolife) and incubation at 37°C for 24 h. The results were expressed as the average (± standard deviation, *n*=4) log colony forming units per gram (log CFU/g) of fruit.

The pH values of fruit samples were measured with a digital pH meter (RL150, Russell pH Cork, Ireland) with a glass electrode (Metrohm AG, Herisau, Switzerland).

### 2.3. DNA extraction of the plate microbiota

After the enumeration of the microbial populations, appropriate countable RBC plates were selected. All the colonies present on the surface of each plate were suspended in 2 ml Ringer buffer solution (Lab M Limited), harvested with a sterile pipette and transferred in a 2 ml vial. The microbial vials were stored by freezing at -80°C supplemented with 50% (v/v) sterile glycerol.

Microbial DNA was extracted as previously described by Hoffman and Winston (1987) with slight modifications. Briefly, 0.3 ml of lysis solution and 0.3 ml of phenol/chloroform were added in the microbial pellets obtained after centrifugation (for the removal of glycerol). The solution was transferred in tubes with 0.3g of glass beads which were then placed in vortex for 4 minutes. The tubes were centrifuged for 2 minutes at 13,000 rpm and the supernatant carefully transferred at 1.5ml tubes. 800 μl of 100% ethanol were added and the tubes were centrifuged at the same conditions. After the centrifugation, 1ml of 70% ethanol was added, the tube was centrifuged again and the supernatant was discarded. The DNA was resuspended in100 μl of TE buffer solution and stored overnight at 4°C. Before further analysis, 1 μl of RNase was added and DNA incubated for 15 minutes at 37°C.

### 2.4. DNA extraction of the pineapple microbiota

Ten grams of each pineapple sample were homogenized with 20 ml Ringer buffer solution (Lab M Limited) in filter Stomacher bag (Interscience, St-Nom, France) for 60 s in the stomacher device. Then, 20 ml of the juice were collected in 50 ml tubes (SARSTEDT AG & Co. KG, Germany) and centrifuged (Heraeus Multifuge 1S-R, Thermo Electron Co.) at 8000 × g for 20 min at 4°C. Since the supernatant was discarded, the microbial pellet was washed with 20ml of distilled-dionized water and centrifuged (Heraeus Fresco 21, Thermo Scientific) again at the same conditions. The cells were rediluted in 1.7ml sterile ultrapure water, transferred in 2 ml eppendorfs (SARSTEDT AG & Co. KG), and centrifuged at 17000 x g for 10 minutes at 4°C. The supernatant was discarded and the microbial cells were stored at -80°C. Microbial DNA was extracted according to the protocol described above for plates.

### 2.5. Barcoding PCRs and Illumina Miseq PCR

Amplicon libraries were constructed following two rounds of PCR amplification. The first amplification of the ITS2 rRNA gene was performed with the primers ITS3 (5’-GCATCGATGAAG AACGCAGC-3’) and ITS4 (5’-3’TCCTCCGCTTWTTGWTWTGC-3’). The final primer concentration used was 10 μM. Forward and reverse primers carried the Illumina 5’CTTTCCCTAC ACGACGCTCTTCCGATCT-3’ and the 5’-GGAGTTCAGACGTGTGCTCTTCCGATCT-3’ tails, respectively. The first round of PCRs was performed with the high-fidelity AccuPrime Taq DNA polymerase system (Invitrogen, Carlsbad, USA) and 5 μL of microbial DNA. The cycling conditions were: 94°C for 1 min, followed by 30 cycles of amplification at 94°C (60 s), 55°C (60 s), and 72°C (60 s), with a final extension step of 10 min at 72°C. The amplicon size, quality, and quantity of the amplified DNA were checked on a DNA1000 chip (Agilent Technologies, Paris, France). Then, the second Miseq PCR was conducted with V3 illumina MiSeq kit as described in Poirier et al. (2018). Raw read sequences were deposited at the Sequence Read Archive under the Bioproject number PRJNA665125 and the accession numbers SAMN16242305 to SAMN16242366.

### 2.6. Quality filtering of reads and taxonomic assignment of Operational Taxonomic Units (OTU)

Raw sequencing reads were imported into the FROGS (Find Rapidly OTUs with Galaxy Solution) pipeline (Escudie et al., 2017) for quality control and assembly into OTU. Roughly, the pipeline was as follow: quality-filtered ITS2 paired-end sequences were merged with VSEARCH v2.15.0 (Rognes et al., 2016) using 0.1 mismatch rate in the overlapped region. Only amplicon with a size above 150 bp and no longer than 500 bp were kept. Merged amplicon sequences were dereplicated and clustered using SWARM v3.0.0 (Mahe et al., 2015) algorithm with a distance threshold of 3. Chimeras were removed with VSEARCH v2.15.0. The resulting sequences were filtered for spurious OTUs by keeping only those with at least 0,01% of relative abundance within the whole dataset (Auer et al., 2017). Taxonomic assignment of OTUs was performed using the UNITE 6.1 ITS2 as reference database (Nilsson et al., 2018 https://unite.ut.ee/) and the Blastn+ algorithm (Camacho et al., 2009).

### 2.7. Analysis of alpha and beta diversity

Fungal diversity was analysed using the R package Phyloseq (McMurdie and Holmes, 2013). OTU abundance was normalized using the median sequencing depth of all samples. Analyses of alpha and beta diversity were carried out using standard or custom Phyloseq command lines.

## 3. Results

### 3.1. Growth of dominant fungal microbiota is temperature dependent

A comparative analysis of total viable mesophilic counts (TVC) and fungal microbial counts on four independent pineapple batches revealed that fungi and mostly yeasts are the main component of the cultivable microbiota over storage (Figure 1). Bacterial population, for both *Pseudomonadaceae* and *Enterobacteriaceae* families commonly found on vegetables and fruits, was therefore very low with no more than 2 log CFU/g throughout storage at all the studied temperatures (data not shown). Moreover, the population of lactic acid bacteria was not detectable with common microbiological analyses due to the dominance of yeasts on MRS agar plates. As it could be expected, the growth of the fungal population was faster at the highest temperatures. The initial level of fungi (mean ± standard deviation, n=4) was 4.69 ± 0.65 log CFU/g, and reached a final average level of 7.36 ± 0.44 and 7.41 ± 0.72 log CFU/g at 8°C and 12°C, respectively. Storage at 4°C revealed more stringent than the three other temperature conditions on growth kinetics. In this case, the fungal population reached only 6.11 ± 0.99 log CFU/g after 230 hours. The microbial growth monitored during dynamic temperature conditions resembled that recorded at 8°C.

**Figure 1.**
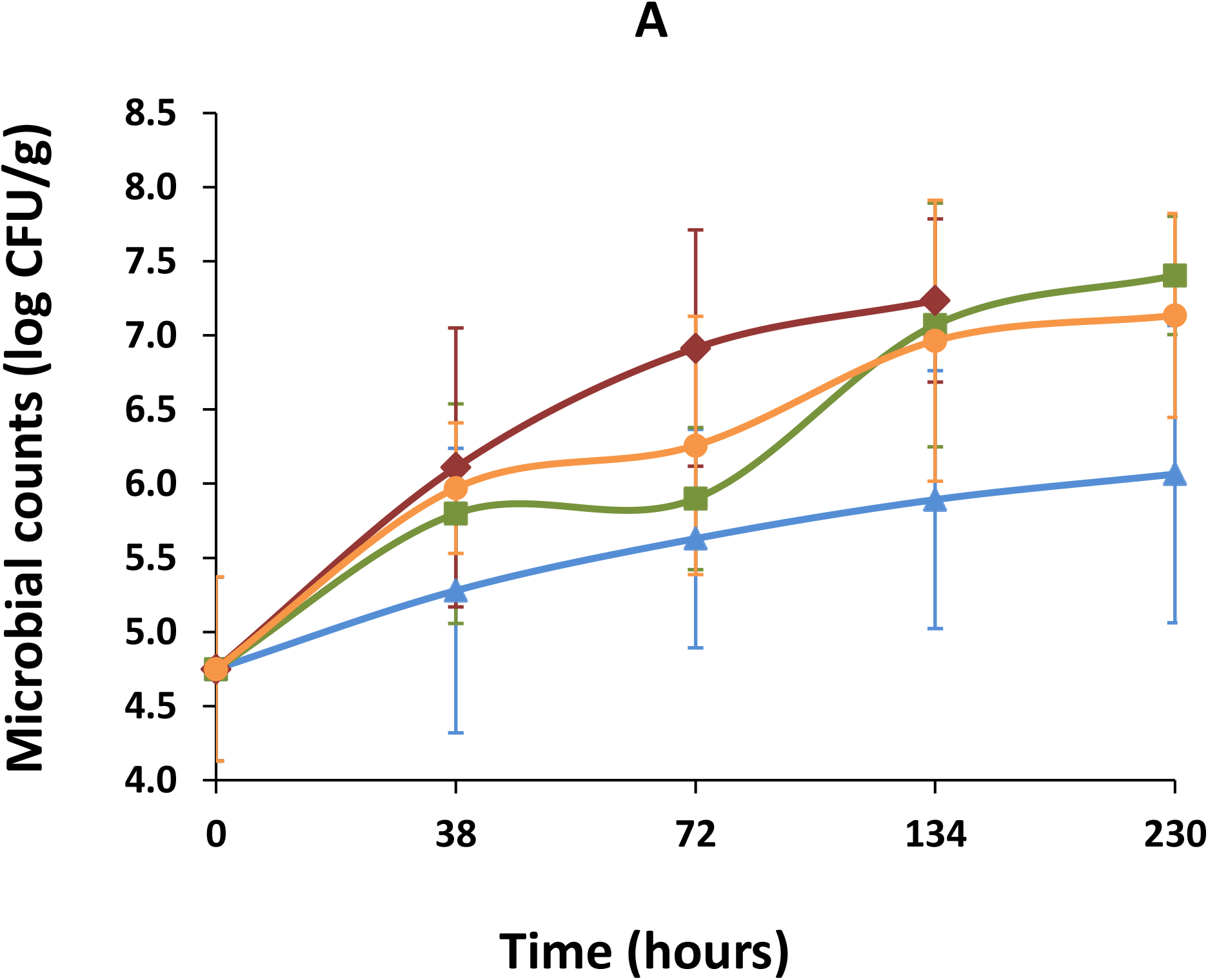
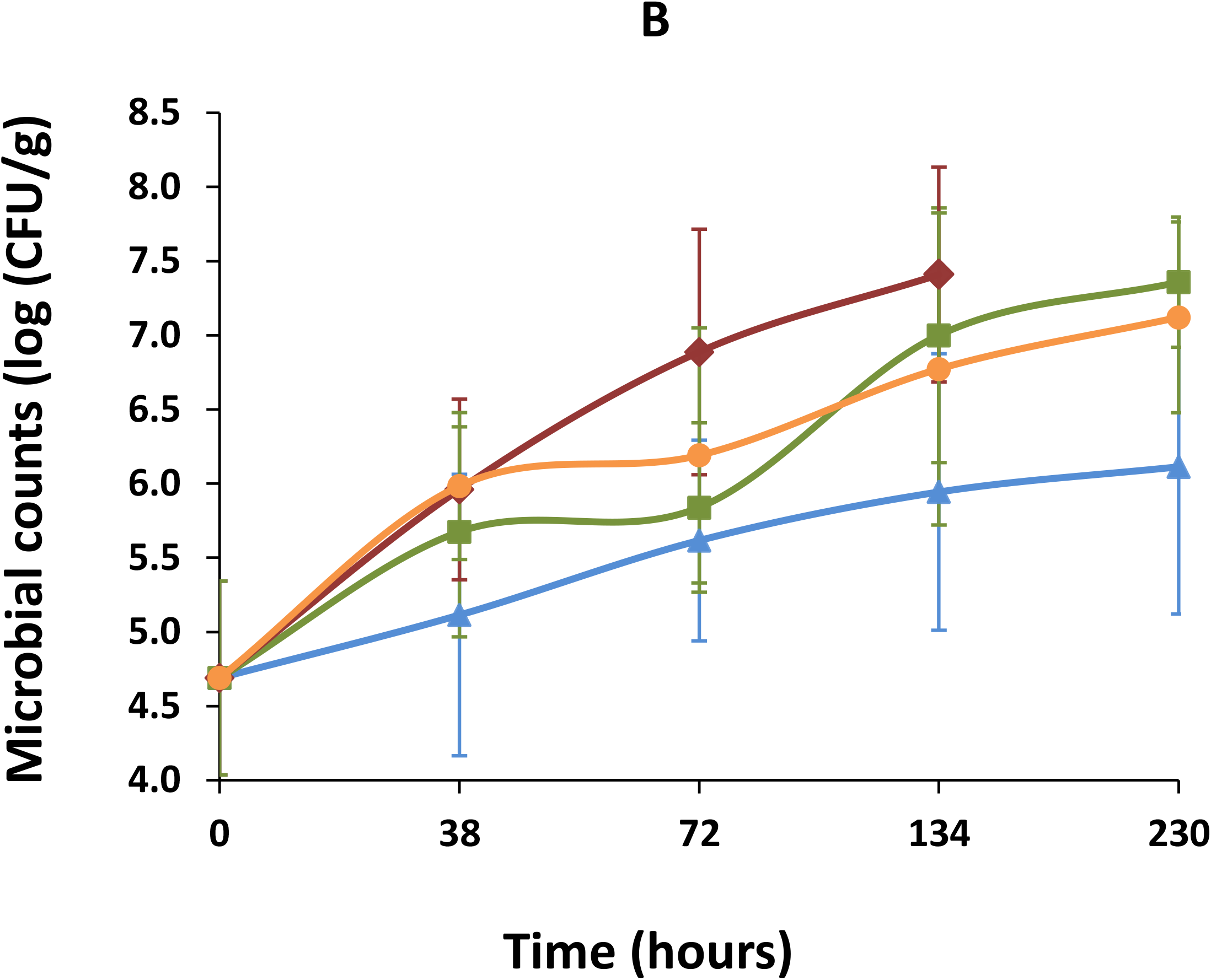
A) TVC populations (mean ± standard deviation, n=4) and B) Fungal populations (mean ± standard deviation, n=4) in RTE pineapple during storage at 4°C, 8°C, 12°C, and dynamic temperature conditions. The blue, green, red and orange lines represent the growth of the microbial populations at 4°C, 8°C, 12°C, and dynamic temperature conditions.

As far as the pH is concerned, no significant differences on pH measurements were found between the different temperatures and during storage (Table 1).

**Table 1.**
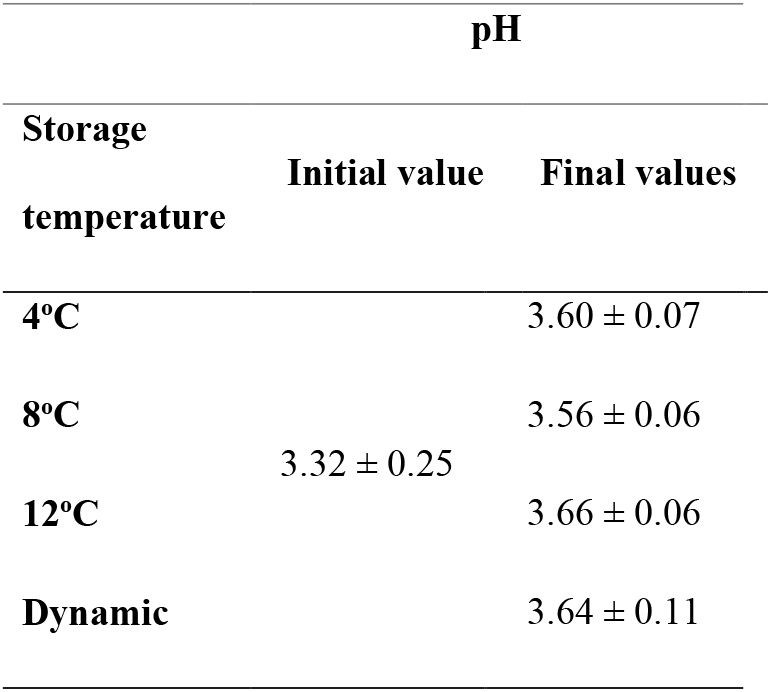
The initial and final pH values of pineapple during storage at different temperatures.

### 3.2. Fungal OTU richness and alpha-diversity is different between pineapple samples and cultivation media

We investigated whether cultivation methods were underestimating the level of fungal population during storage. We compared the fungal diversity by ITS2 amplicon sequencing between DNA extracted directly from pineapple samples or from the fungal population that grew on agar plates. As shown in Table 2, the fungal OTU richness was significantly lower on plates.

**Table 2.** Fungal OTU richness (merged at different taxonomic levels) between food pineapple samples and agar plate samples.

|  | Number of genera | Number of species | Number of OTU |
| --- | --- | --- | --- |
| **Pineapple samples** | 22 | 33 | 47 |
| **Plate samples** | 18 | 25 | 39 |

Figure 2 shows how the 33 fungal species detected by non-cultural metagenetic analysis could be detected and quantified from the microbiota recovered from agar plates. In general, detection of most species from Basidiomycota phylum was unsuccessful in comparison to species from Ascomycota phylum. In addition, detection of *Fusarium* species was also strongly biased on plates, in particular *Fusarium circinatum* was highly abundant in pineapple samples compared to plates. Therefore, to avoid any bias in our analysis, only data from pineapple samples were used further.

**Figure 2.**
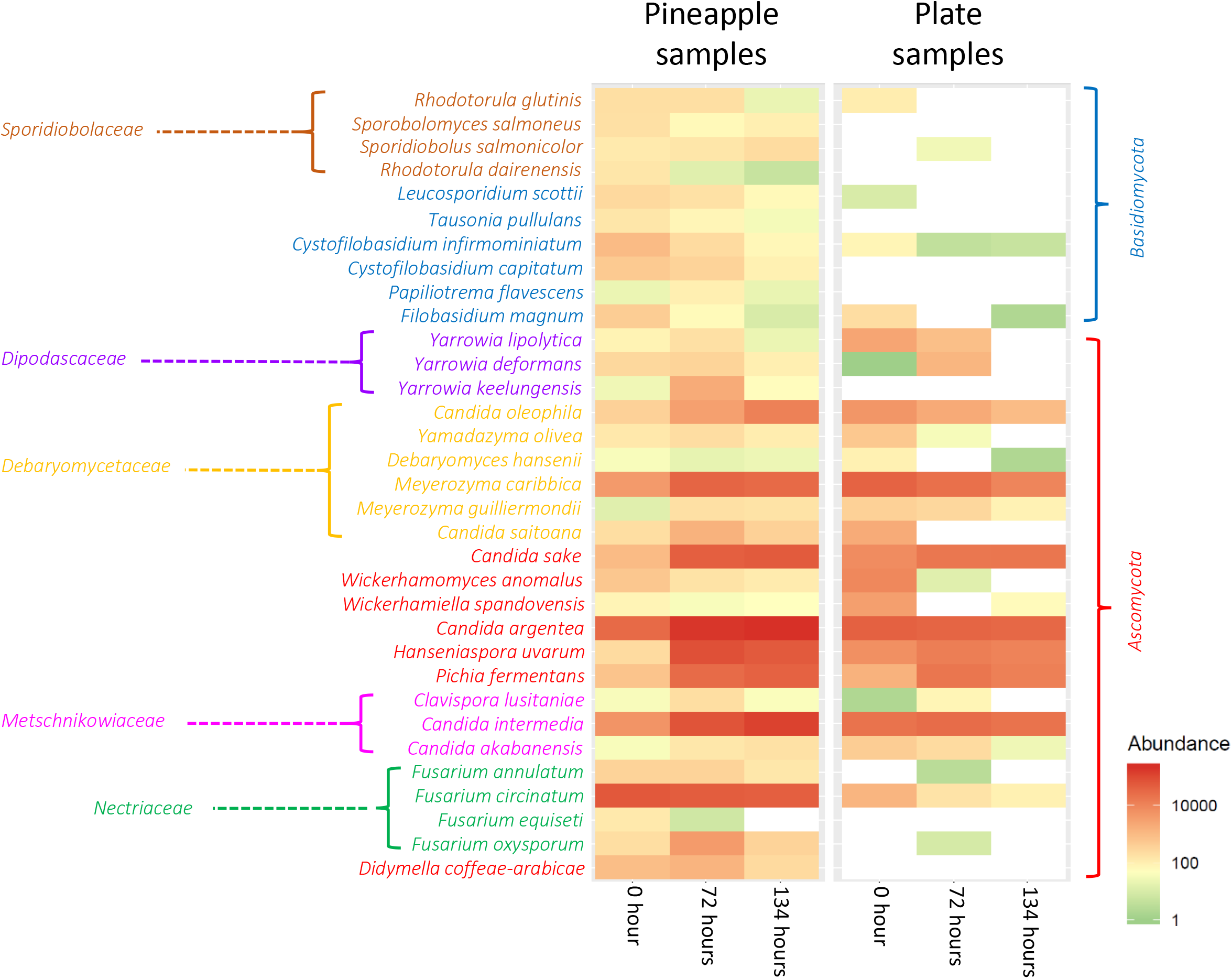
Heatmap showing the comparison between fungal species relative abundances detected by non-cultural methods (Pineapple samples) and by cultural methods (Plate samples). Each column shows the average relative abundance of the 33 species after the various samples from the four pineapple batches and for the different storage temperature were merged. Main fungal families and phyla are depicted by bracklets.

### 3.3. The effect of temperature and time of storage on fungal diversity

The effect of different temperatures and storage times was analysed on fungal richness after merging OTUs at species-level (Figure 3). The species’ richness was significantly higher in pineapple samples stored at 4°C (p<0.01). The samples stored at dynamic conditions and 8°C followed, while the samples stored at the highest temperature (12°C) had the lowest number of species. The fungal richness was also comparable for the different storage time (p<0.01). At the beginning of the storage (zero time), the diversity was higher compared to all the other storage times. Although the species richness decreased over time, the species’ number at 230 hours did not follow the same declining course. This observation is not unexpected, since the samples at 230 hours come exclusively from storage at 4°C. The corresponding samples (230h) stored at 8°C and dynamic conditions were not successfully sequenced.

**Figure 3.**
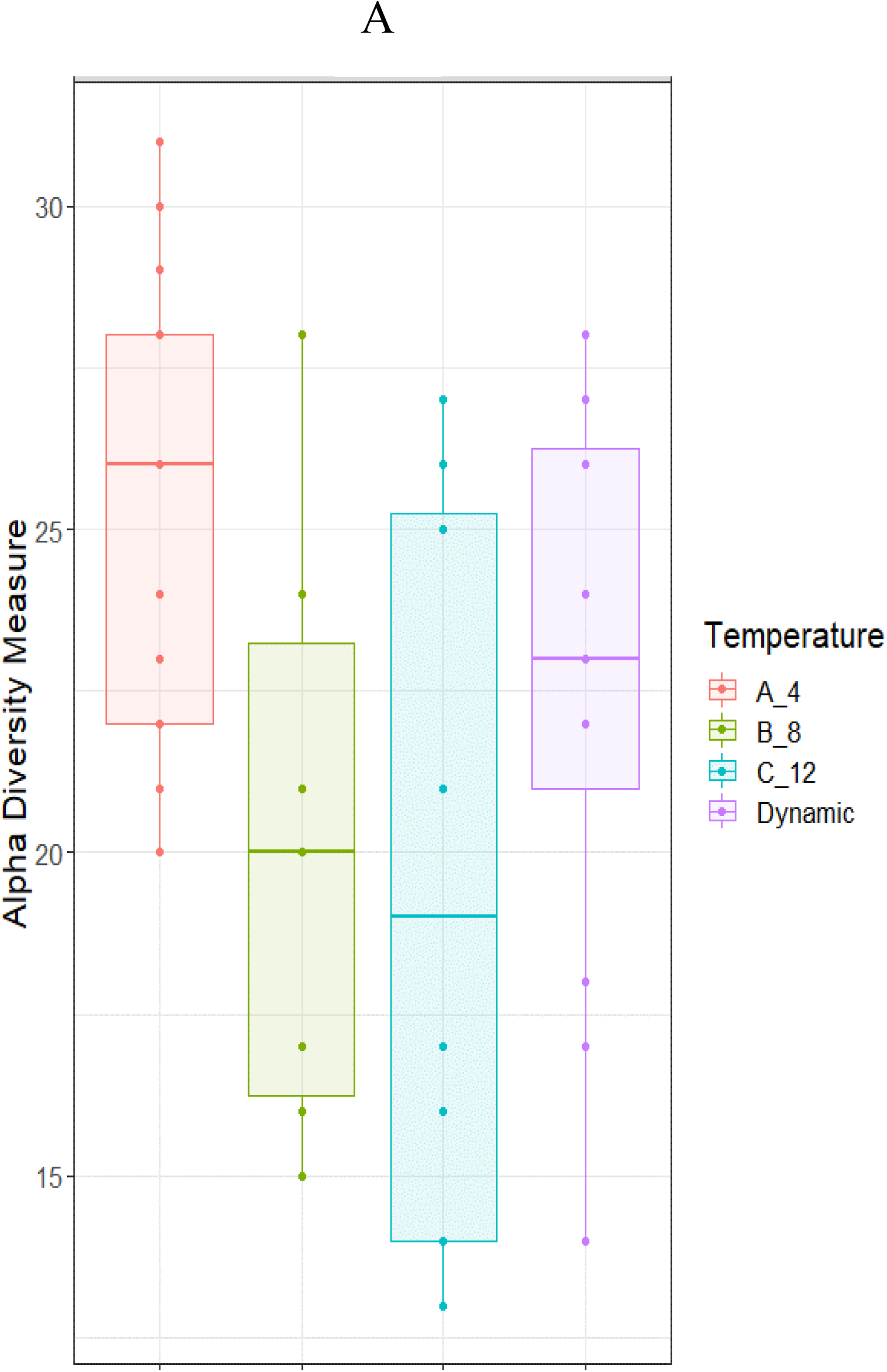
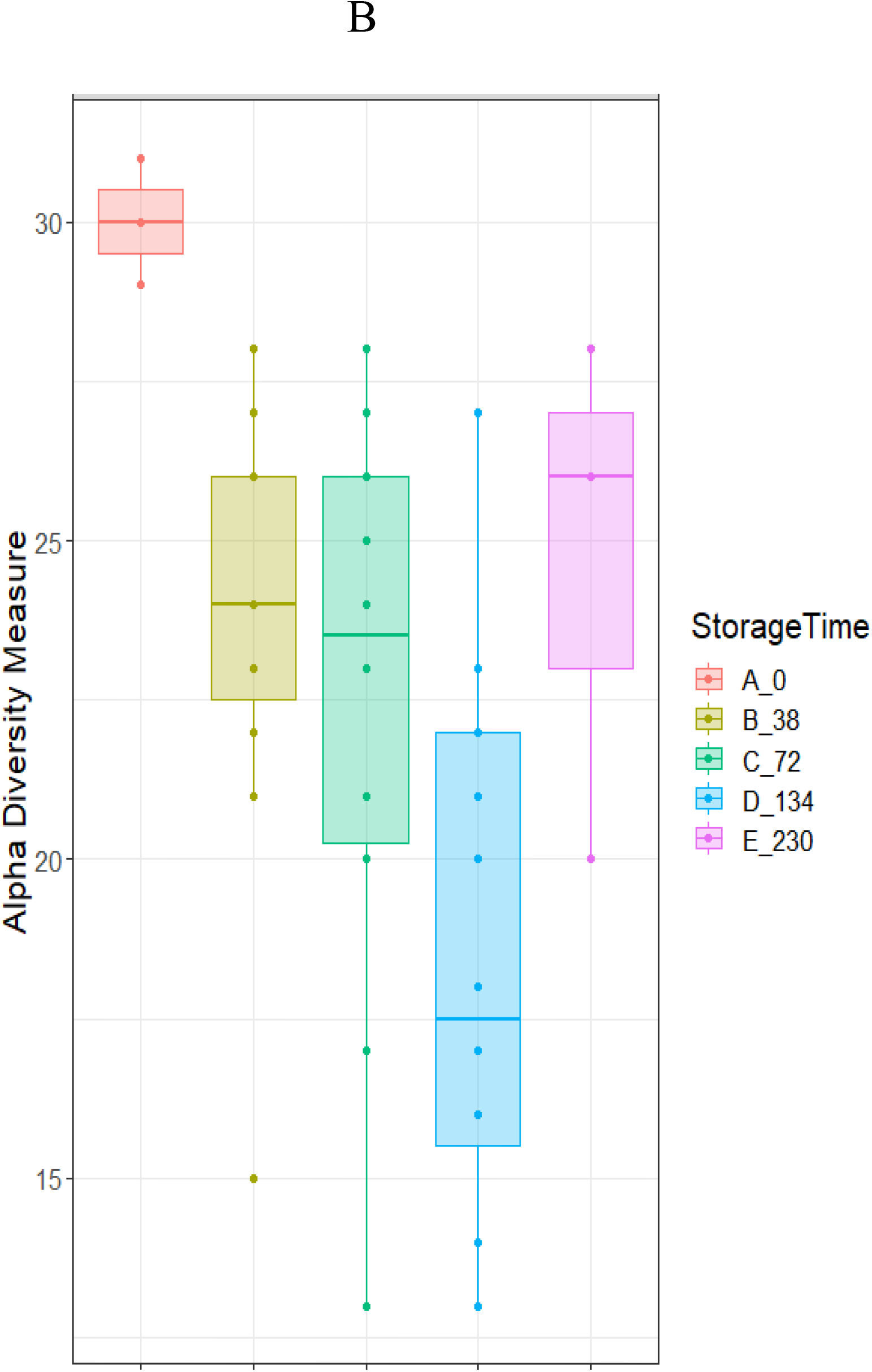
Fungal richness in RTE pineapple samples. The box plot shows the number of species in samples of different temperatures (A) and time (B) of storage. The boxes represent the interquartile range between the first and third quartiles and the vertical line inside the boxes is the median obtained from the samples analysed per condition.

We also performed Non-Metric Multidimensional Scaling (NMDS) on Bray-Curtis distances to statistically compare the fungal diversity within the samples of different temperature and time of storage. In all cases (data not shown), communities recovered from a given temperature or storage time did not clustered together on the factorial plane. On the other hand, there were a discrete clustering among samples of the different batches of pineapple, presented in Figure 4. The fungal diversity of most of the samples from batch P1 and P2 differs with that of batches P3 and P4 together.

**Figure 4.**
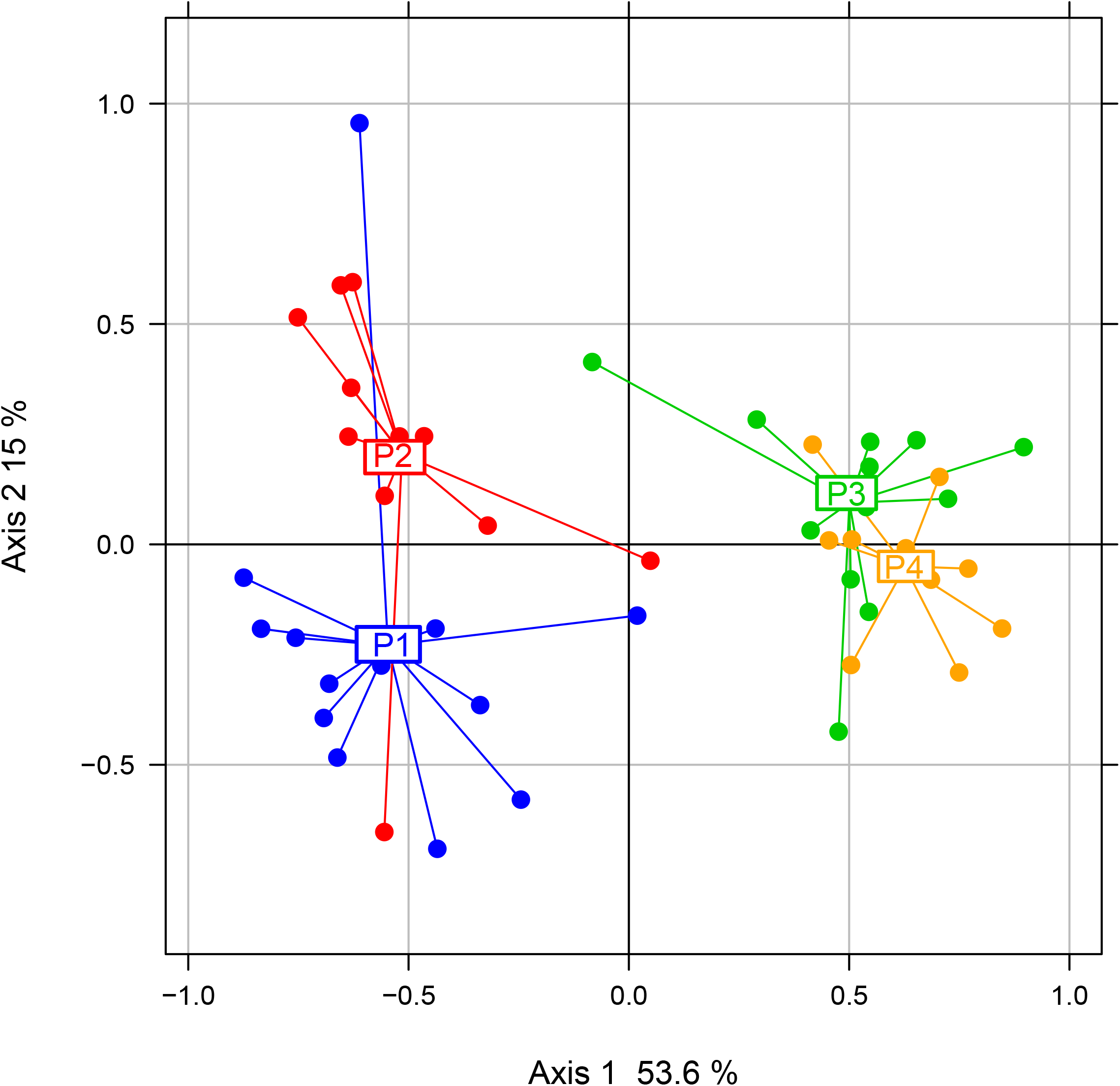
Non-Metric Multidimensional Scaling (NMDS) based on Bray-Curtis distances among fungal communities of the four pineapple batches. P1, P2, P3 and P4 correspond to batch 1, batch 2, batch 3 and batch 4.

The differences on fungal diversity between the four batches are presented in Figure 5, where the composition plot of relative abundances is illustrated according to Bray-Curtis hierarchical clustering of pineapple samples. In Figure 6 is also presented a phylogenetic tree of the different fungal species based on the ITS2 sequences.

**Figure 5.**
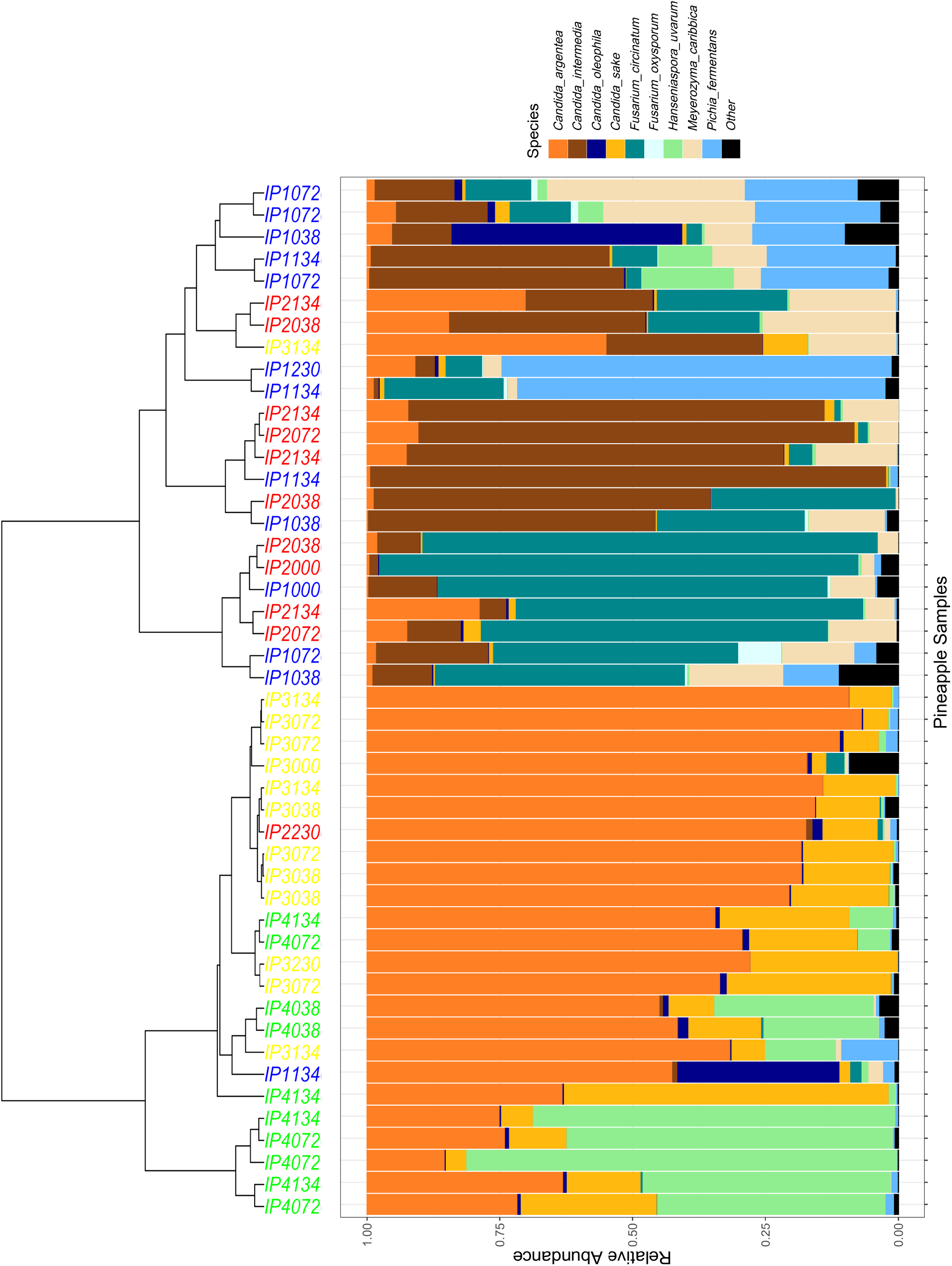
Composition plot showing the relative abundances of the nine main Ascomycota species found in Pineapples samples. On the top: hierarchical clustering of batches samples according to Bray-Curtis distance and ward algorithm (blue for P1, red for P2, yellow for P3 and green for P4 as shown in Figure 4).

**Figure 6.**
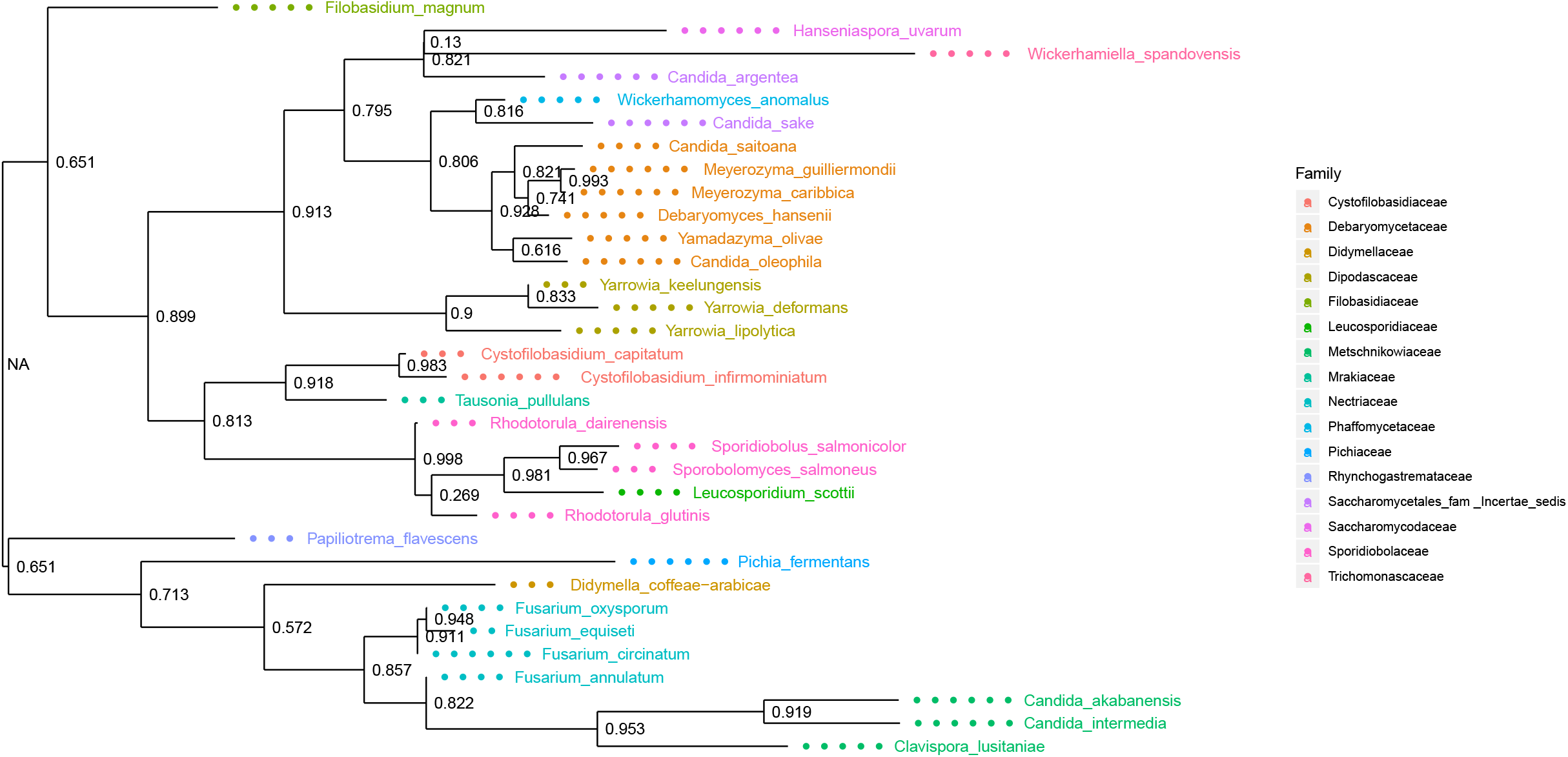
Neighbour-Joining phylogenetic tree of the different species based on the ITS2 sequences of pineapple samples. Bootstrap values are indicated on the main nodes.

Pineapple samples from batches P3 and P4 had a quite similar fungal community dominated by two phylogenetically related species *Candida argentea, and Candida sake* from the candidate family *Saccharomycetales_incertae_sedis*. Most samples from Batches P4 can be distinguish from those of batch P3 with the presence of *Hanseniaspora uvarum*, which is also closely related phylogenetically to the aforementioned *Candida*. Pineapple samples from batches P1 and P2, displayed a set of completely different fungal communities dominated by species from the phylogenetically related *Nectriaceae* (*Fusarium*) and *Metschnikowiaceae* (*Clavispora/Candida*) families. *Candida intermedia* and *Fusarium circinatum* were the most abundant species in samples of batch P2, whereas some samples from batch P1 showed higher level of diversity with *Pichia fermentans* (*Pichiaceae*) and *Meyerozyma caribbica* (*Debaryomecytaceae*).

At batch level, the effect of storage time and temperature varied across the different fungal species. Interestingly, the initial composition of pineapple microbiota had a great impact on the evolution of spoilage at different temperatures and storage time. Therefore, situations varied from one batch to another.

Starting with Batch 1 (Figure 7), we observed that both temperature and storage time drove a strong change on the fungal community structure. Temperature was the most influent parameter separating samples stored at 4°C from those stored at 12°C, with in both cases, a gradual change over longer storage time. Samples stored at intermediate temperature of 8°C, clustered at intermediate positions between samples stored at 4°C or 12°C. Only samples stored with a dynamic sequence of temperature scattered randomly with no logical order. Among the most striking changes, we observed that *F. circinatum* had high abundance (>75%) at zero time of storage, while following the next stages of storage *C. intermedia* or *P. fermentans* finally dominated according to the temperature.

**Figure 7.**
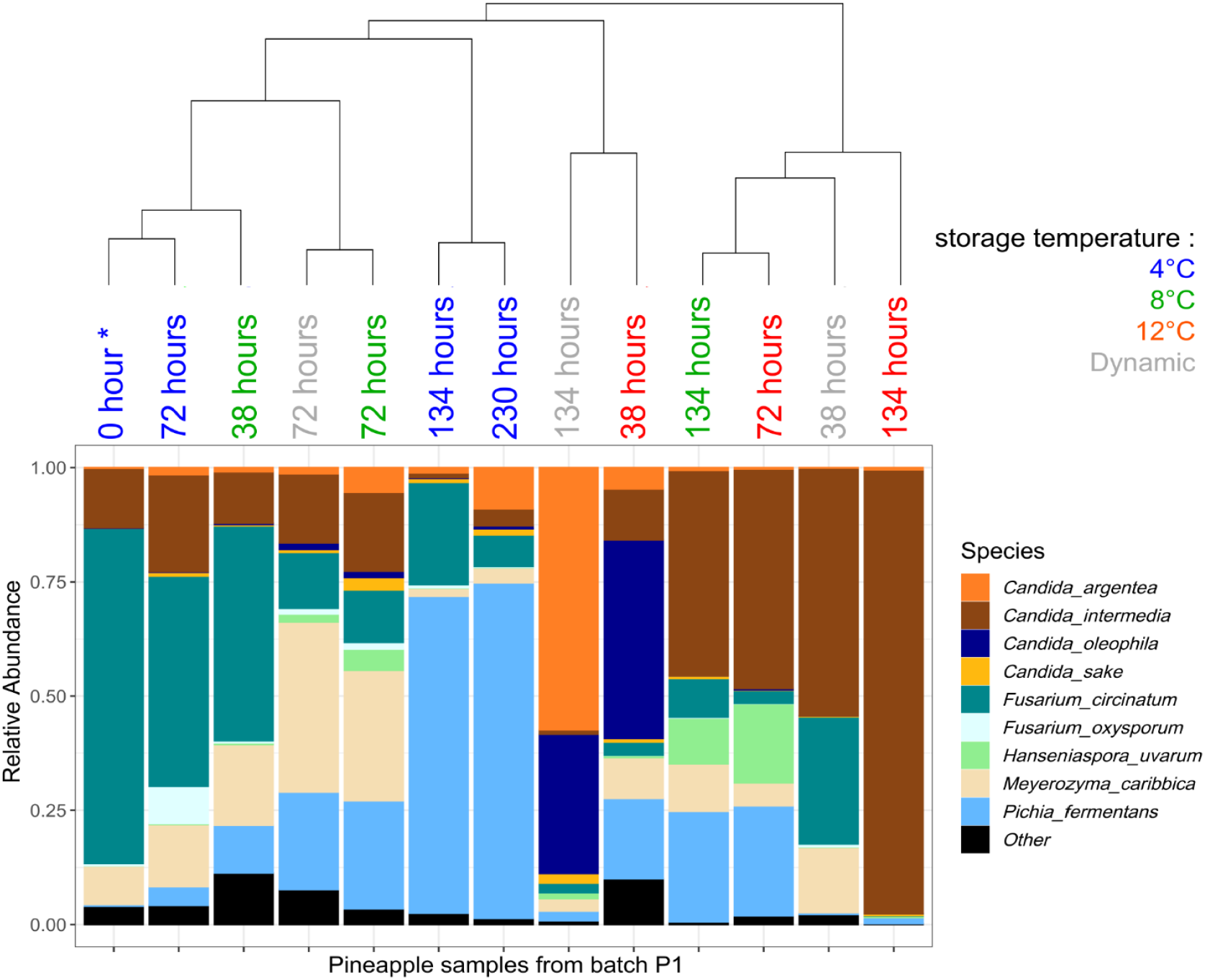
Impact of storage time and temperature on fungal species composition of pineapple samples from Batch P1. Samples are ordered from left to right according to Bray-Curtis distance. The asterisk (0 hour) indicates the initial analysis before packaging and storage at any other conditions.

Specifically, *C. intermedia* succeeded to dominate to samples of 12°C and 8°C from the early stages of storage, while *P. fermentans* in samples of 4°C at the final stages (134 and 230 h). On the other hand, *M. carribica* was able to dominate only at the middle of storage (72 h) for 8°C and dynamic conditions. Concerning batch P2 (Figure 8), a trend similar to that observed in batch P1 can be drawn with, at time zero, a large dominance of *F. circinatum* which was progressively replaced by *C. intermedia* at high storage temperatures and by *C. argentea* at 4°C. As shown in both Figure 4 and Figure 5, the fungal communities from samples of batches P3 and P4 were not affected significantly by temperature and storage time. Unlike pineapple samples from batches P1 and P2, the fungal community of batch P3 was covered by the great dominance of *C. argentea* throughout the storage at all temperatures (data not shown). *C. sake* was not affected by temperature or time in P3 but also in P4. However, in the case of samples from batch P4, we noticed a slight impact of temperature and storage time on *H. uvarum* and *C. argentea*. The former became more prevalent at 8 and 12°C in the middle of storage, while the latter prevailed at 4°C throughout storage and at 12°C and dynamic conditions at the early stages of storage.

**Figure 8.**
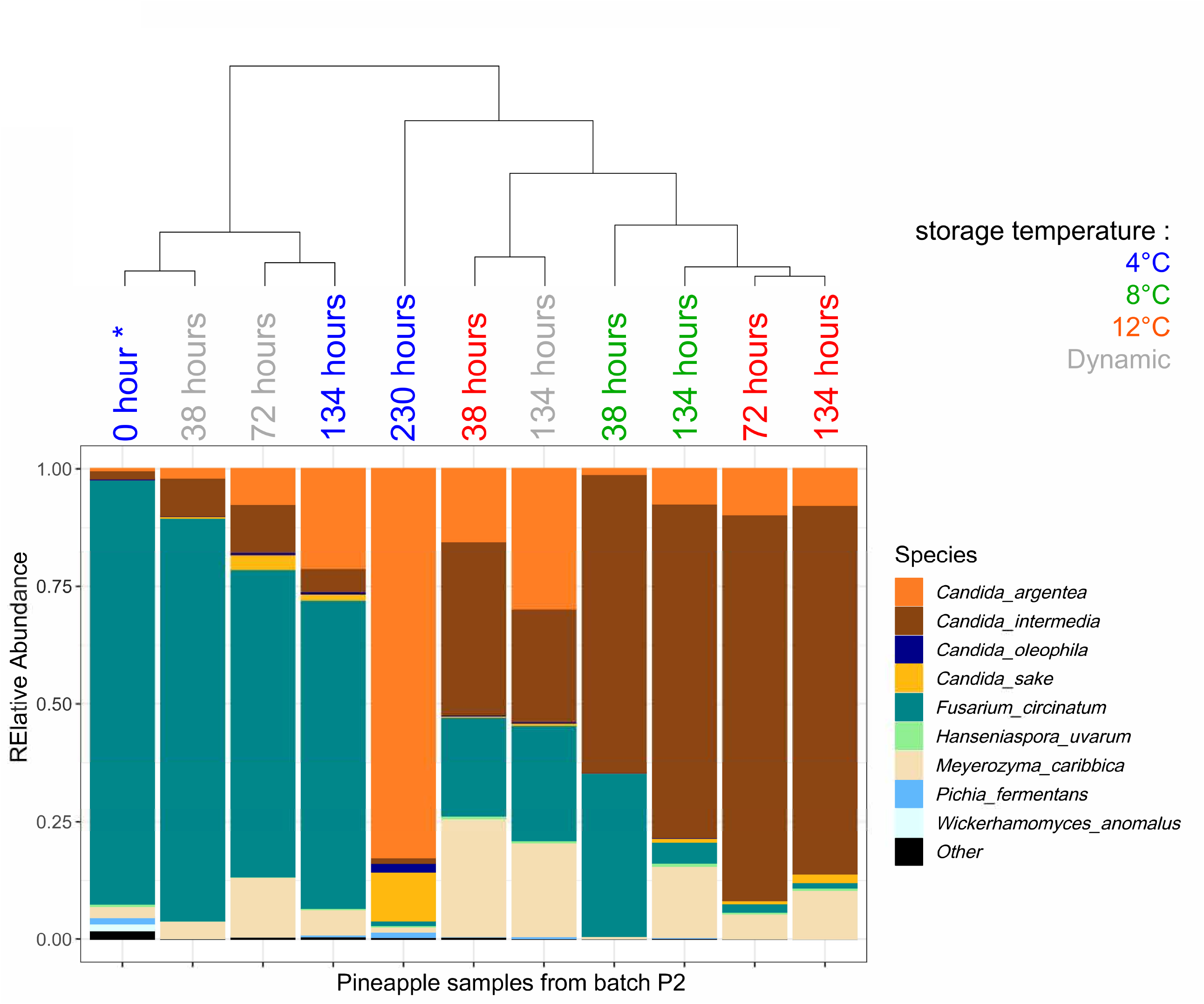
Impact of storage time and temperature on fungal species composition of pineapple samples from Batch P2. Samples are ordered from left to right according to Bray-Curtis distance. The asterisk (0 hour) indicates the initial analysis before packaging and storage at any other conditions.

## 4. Discussion

The aim of the present study was to characterize the microbial community involved in the spoilage of RTE pineapple and determine the changes of the diversity when stored under different temperatures, using a metagenetic approach.

Our results demonstrated that fungi and mainly yeasts were the predominant spoilage microorganisms found in RTE pineapple. Not surprisingly, the high yeast population is likely attributed to the high level of sugars and the low pH of pineapple, which is ideal for their growth (Leneveu-Jenvrin et al., 2020; Da Cruz Almeida et al., 2018; Maciel et al., 2013). Previous works, recorded similar levels of mesophilic populations and fungi in pineapple (Leneveu-Jevrin et al., 2020; Di Egidio et al., 2009; Montero-Calderon et al., 2008; Tournas et al., 2006). Two of these studies also pointed out significant differences on the initial and final microbial values depending on the different batches (Leneveu-Jevrin et al., 2020; Tournas et al., 2006).

Although cultural methods have been extensively be used in food microbiology, they are also considered extremely biased in their ability to capture the microbial diversity of complex environments (Cao et al., 2017; Edet et al., 2017; Ercolini et al., 2013). Consequently, we firstly proceeded to a comparative analysis of the culture-dependent and culture-independent characterization of fungal community of pineapple. We were able to demonstrate that a whole phylum (Basidiomycota) was hardly detected in plates, while one of the most abundant species detected in one batch (*Fusarium circinatum*) was unsuccessfully represented in plates’ microbiota. To our opinion, this important finding indicates that non-cultural analytical methods should be advised for further microbiota analysis of pineapple.

The effect of temperature and storage time on fungal diversity of pineapple samples was further analysed. Both factors had a significant impact; specifically, species richness decreased over storage time and when the temperature reached higher levels. However, when we compared the fungal diversity within the samples of different temperature and time of storage based on Bray-Curtis distances, no radical clear clustering could be evidenced according to the studied factors. On the other hand, the analysis revealed a significant batch effect on fungal diversity and composition. Leneveu-Jenvrin et al. (2020) also observed that the pineapple communities clustered mostly according to batches and not storage time. The high biological variability is a common observation in plant origin products due to the strong impact of various factors such as cultivar, geographical region, and agricultural practices (Leff and Fierer et al., 2013).

In the present work, the two batches (P3 and P4) had similar composition, since the phylogenetically related *C. argentea* and *H. uvarum* (only in batch P4) or *C. sake* were the dominant species. On the other hand, most samples from P1 and P2 were dominated by the species *F. circinatum* or *C. intermedia* which are also phylogenetically related to each other. Moreover, some samples of P1 showed higher level of diversity, since *P. fermentans* and *M. caribbica* were present in some samples in high abundances. There are few previous studies (not NGS studies) concerning the fungal composition of pineapple, mainly based on culture-dependent techniques. Tournas et al. (2006) detected *Schwanniomyces polymorphus* (formerly known *Debaryomyces polymorphus), Candida pulcherrima, Pichia* spp. and in low abundances *Penicillium* spp., while Di Cagno et al. (2010) found *Meyerozyma guilliermondii* (formerly known as *Pichia. guilliermondii)* as the only species grown on plates. Chanprasartsuk et al. (2010) identified *M. guilliermondii* and *H. uvarum* as the main yeasts of fresh pineapple juices from different locations and countries. These two species were characterized both by culture-dependent and independent techniques (using DGGE). Zhang et al. (2014) used *Candida argentea, Candida sake* and *Meyerozyma caribbica* isolates for the investigation of the headspace oxygen level in shelf life of pineapple, since they were previously isolated from spoiled commercial fresh-cut pineapple. Ibrahim et al. (2017) identified *Fusarium proliferatum, Fusarium verticillioides, Fusarium sacchari* and *Fusarium* sp. in diseased pineapple tissues. Some of the *Fusarium* sp. isolates appeared to be phylogenetically related to *F. circinatum.* Recently, Lima et al. (2019) found that in tropical fruit based ice-creams (including pineapple based) the predominant species were *Candida intermedia, Torulaspora delbrueckii, Candida parapsilosis, Clavispora lusitaniae, Saccharomyces cerevisiae* and *Pichia kudriavzevii.* It is obvious that there are differences in composition between the various studies. As it was mentioned above, these differences could be attributed to various pre- and pro-harvest environmental factors, agricultural practices, but also plant genotype. Additionally, there is a low species diversity in previous studies which may be related to the biases of culture-dependent techniques (as we discussed above) or the limitations of conventional genomic methods (Subasinghe et al., 2019).

In the light of this scientific literature and by comparing them with the data obtained in our work, it is plain that pineapple may harbour a vast variety of fungal microbiota. However, our study provide additional information on how these different microbiotas may behave towards storage conditions. A thorough observation at batch-level reveals various conclusions about the impact of temperature and storage time. Specifically, the batches P1 and P2 were characterized by the great dominance of *F. circinatum*. Multiple species of *Fusarium* are associated with fruit rot and leaf spot diseases of fruits and especially pineapple (Jacobs et al., 2010). However, at the later stages of spoilage in batch P1, *Pichia fermentans* prevailed at 4°C, while *C. intermedia* prevailed at 8 and 12°C. Both *P. fermentans* and *C. intermedia* have been studied with great potential in the control of phytopathogenic molds. Rosa-Magri et al. (2011) identified *C. intermedia* as one of the yeasts with biocontrol activity against *Colletotrichum sublineolum* and *Colletotrichum graminicola*. Giobbe et al. (2007) investigated the dual nature of a strain of *P. fermentans* which controls brown rot on apple fruit, but becomes a destructive pathogen when applied to peach fruit. Consequently, it could be postulated that these two yeasts could possibly play a competitive role in supressing the growth of *F. circinatum* and according to the temperature, one of the two closely related yeasts is able to dominate. The same trend was followed in batch P2, but *C. argentea* prevailed finally at 4°C. On the contrary, when *C. argentea* was present in great dominance (batch P3) at first place there was no significant impact of temperature and time. The conclusion is differentiated (batch P4) when *H. uvarum* was initially present together with *C. argentea* as the second most dominant species. The two closely related species seem to be affected by the temperature and time in an opposite way, but in a lesser extent. Consequently, the progress of spoilage firstly depends on the initial composition and secondly is determined by the effect of temperature and time.

## 5. Conclusions

The current study significantly contributes to the understanding of the fungal community associated to fresh-cut and RTE pineapple. It is the first time that the impact of temperature and storage time on fungal diversity is being studied for a fresh tropical fruit product. The results demonstrated that the different batches of pineapple show great variability on fungal composition. The initial composition constitute an important factor on spoilage progress. Depending on the initial prevalent fungal species, the impact of temperature and storage time varies. It is obvious that fresh pineapple products are a very complex and unpredictable ecological niche where the specific spoilage species may have a totally different response to the changes of important environmental factors used for storage and which are important to assess the shelf-life of these RTE fruit. Consequently, further and thorough research are necessary in order to unravel how the various environmental factors of pineapple production drive the initial pineapple microbial composition. In this view, a large-scale analysis from various production facilities must be advised.

## 6. Acknowledgments

We thank the MIGALE bioinformatics platform at INRAE (http://migale.inrae.fr) for providing computational resources and data storage. We also thank the INRAE@BRIDGe platform for carrying out the MiSeq sequencing runs.

This research did not receive any specific grant from funding agencies in the public, commercial, or not-for-profit sectors.

